# James-Stein estimator improves accuracy and sample efficiency in human kinematic and metabolic data

**DOI:** 10.1101/2024.10.07.616339

**Authors:** Aya Alwan, Manoj Srinivasan

## Abstract

Human biomechanical data are often accompanied with measurement noise and behavioral variability. Errors due to such noise and variability are usually exaggerated by fewer trials or shorter trial durations, and could be reduced using more trials or longer trial durations. Speeding up such data collection by lowering number of trials or trial duration, while improving the accuracy of statistical estimates, would be of particular interest in wearable robotics applications and when the human population studied is vulnerable (e.g., the elderly). Here, we propose the use of the James-Stein estimator (JSE) to improve statistical estimates with a given amount of data, or reduce the amount of data needed for a given accuracy. The JSE is a shrinkage estimator that produces a uniform reduction in the summed squared errors when compared with the more familiar maximum likelihood estimator (MLE), simple averages, or other least squares regressions. When data from multiple human participants are available, an individual participant’s JSE can improve upon MLE by incorporating information from all participants, improving overall estimation accuracy on average. Here, we apply the JSE to multiple time-series of kinematic and metabolic data from the following parameter estimation problems: foot placement control during level walking, energy expenditure during circle walking, and energy expenditure during resting. We show that the resulting estimates improve accuracy — that is, the James-Stein estimates have lower summed squared error from the ‘true’ value compared with more conventional estimates.

## 1 Introduction

Measured biomechanical data are often accompanied with sources of error such as incomplete or missing data, measurement noise, and high human variability [1–5]. The effect of measurement noise and movement variability can be sometimes mitigated using signal processing tools such as band-pass filters or even simple averaging [2, 6]. Such defense against noise and variability is aided by having larger sample sizes: conversely, insufficient sample sizes can result in low statistical power and poor replicability [7]. Here, we present the use of an underutilized statistical approach, the James-Stein estimator, heretofore not used for biomechanical data — to enhance accuracy of estimated biomechanical variables especially when the sample size is limited [8–11].

Our focus here is on time-series data, where an analog of sample size is the duration for which the time-series data is collected: usually, longer duration trials provide higher accuracy in parameter estimation [4, 12–14]. For example, it is conventional to average over two to three minutes of steady state metabolic data to obtain an acceptable mean metabolic energy rate (i.e. metabolic cost) value during walking or other exercise [15–20]. Similarly, authors have characterized the minimum number of walking steps required to estimate step kinematic variability during treadmill-walking [21], how sample size and number of steps affects running biomechanical estimates [13], and how quantities derived from step-to-step locomotor variability related to stability and control increase in accuracy with the number of steps [4, 5, 12]. However, the number of steps or the duration of walking during an experiment may be limited due to functional impairment in populations such as the elderly, amputees with prosthesis or exoskeletons, or individuals with musculoskeletal disorders. In such cases, methods to reduce trial duraction while retaining estimation accuracy are of particular interest. Similarly, reducing the trial duration may be useful in human in the loop optimization [22, 23] of wearable devices such as robotic exoskeletons and prostheses, where a large number of trials may need to be performed. Here, we show how the James-Stein estimator allows such reduction of trial duration for a given accuracy of the statistical estimate.

The simplest way to obtain a statistical estimate of a quantity from many samples or long trial durations is via averaging. Other estimation methods include linear and nonlinear regression. Averaging, linear regression, or nonlinear regression are special cases of a broader class of parameter estimation methods called ‘maximum likelihood estimation’ (MLE): given a model of the noise, maximum likelihood estimation computes the parameter values that makes the observed data have the maximum likelihood. MLE methods are popular due to their satisfying the properties of asymptotic normality, consistency, and efficiency [24]. Despite the popularity of MLE (especially its aforementioned special cases), Stein [8, 9] showed that in some situations, there exists a better estimator than the maximum likelihood estimator with lower summed squared error, so MLE is considered ‘inadmissible’. The James-Stein estimator [8–11] (JSE) is one such better estimator.

JSE was popularized in the 1970s by a baseball example [10, 25]. Researchers first computed the batting averages for each player over the first 45 at bats (Fig. 1a), that is, using just the initial few games of the baseball season. These averages (‘MLEs’) provide an inaccurate estimate of the true batting average over an entire season, which typically has over 450 at bats (median). Then, the researchers applied the James-Stein estimator correction for each player, based on all the players’ initial season averages, resulting in new JSE-based estimates. Remarkably, these JSEs were better estimates of the players’ full season averages — even though the JSEs are still based on just the initial part of the season and do not use the full season data. Specifically, the JSEs had lower summed squared error compared with the MLEs when averaged across all participants. Our goal is to examine whether such a result would be true in a variety of biomechanical time-series data, improving the accuracy of shorter duration trials by combining data from multiple participants via JSE (Fig. 1b).

**Fig. 1.**
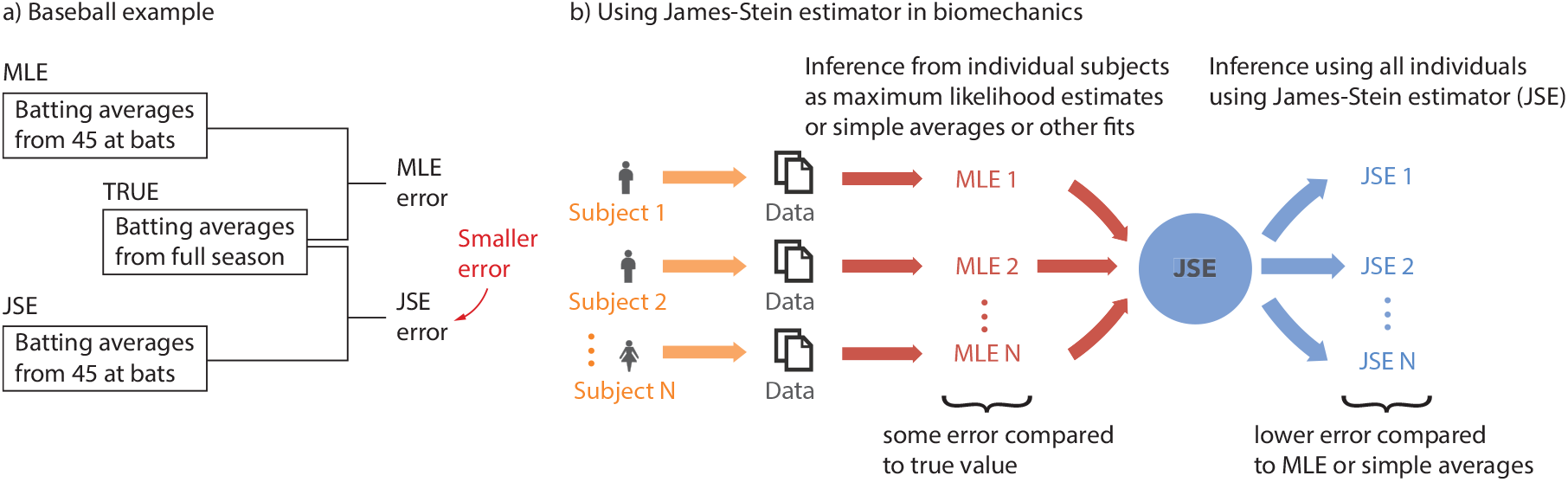
The potential of the James-Stein Estimator. a) Famous baseball example illustrating the potential of using JSE to reduce error, compared with simple averages or MLE [10, 25]. b) Using James-Stein estimator in biomechanics. Statistical estimates of parameters can be obtained for individual participants using averaging, regression, or maximum likelihood estimation (MLE). For limited sample sizes or trial durations per participant, MLEs have some error compared with the true value. The JSE uses information from all the MLEs to produce a new parameter estimate for each participant, which on average will have lower error than those from MLEs.

James-Stein estimation sits within a broader class of meta-analysis-related methods, usually used to obtain a common mean value from multiple studies, but sometimes also used improve the estimates of individual means when appropriate: related methods include Bayesian shrinkage methods, empirical Bayes estimators, isotonic regression, pretest estimators, and other regularized estimators using ridge or lasso regularization [26–29]. Such estimators have been shown to be useful recently in a variety of biomedical contexts involving diabetes [27], cancer [27, 30–32], Alzheimer’s disease [27], COVID-19 [30], Creutzfeld-Jacob disease [28], spectroscopy [33], genomics [34, 35], blood pressure [36], dermal issues [37], body fat [38], and neural recordings [39–41]. Here, we focus on James-Stein estimation for its computational simplicity and target biomechanical estimation problems not previously investigated with such techniques.

In this manuscript, we apply JSE to the following human time-series datasets and related estimation problems: (1) estimating foot placement control during walking derived from from pelvis and foot kinematic data [5]; (2) estimating the steady state metabolic energy rate (*Ė*_walk_) during walking in a circle derived from metabolic time-series [18]; and (3) estimating the mean resting metabolic energy rate (*Ė*_rest_) during quiet sitting [17, 18, 42, 43]. We hypothesize that we can achieve significant improvement in estimation accuracy when utilizing JSE compared with MLE (Fig. 1) and find that JSE does accomplish such error reduction.

## 2 Methods

We first describe how the JSE correction is done if we have an initial estimate and how we compare the results with the ‘true’ value of the estimate. We then describe the experiments and datasets that we consider for JSE, the biomechanical quantities estimated with each dataset, and how these quantities are estimated. All protocols were approved by the Ohio State University IRB and participants took part in the experiments with informed consent.

### 2.1 Computing JSE for better person-specific estimates

Say we have *k* individual participants and we have computed the maximum likelihood estimates (MLEs) *y*_*i*_ (*i* = 1 … *k*) of some quantity for each of these participants. These MLEs could be could be of any quantity of interest and could be obtained by linear regression, simple averaging, or some other nonlinear fitting process, as described more in section 2.2. The corresponding JSE *z*_*i*_ for each participant is determined by shrinking the individual MLEs *y*_*i*_ toward the global MLE average 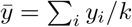 by a ‘shrinkage factor’ *c* using the following remarkably simple equation [8, 10]:

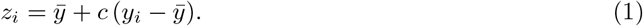

The shrinkage factor *c* in James-Stein estimation is evaluated as:

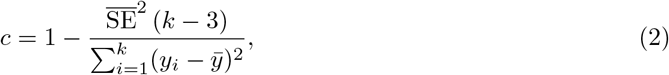

where 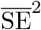 is the variance of the individual MLEs, thus an estimate of errors in the MLE [6]. If the individual MLEs have unequal variances, 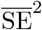, the mean of these variances is used. A shrinkage factor *c* of 1 means that the JSE for an individual is the same as its individual MLE and a *c* value of 0 means that the JSE shrunk the values by 100% toward the global MLE average, so that now all the JSE’s are equal to the global MLE average 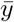 [8, 10].

To assess the performance of the JSE in reducing estimation error, we computed summed squared error of the MLEs *y*_*i*_ and JSEs *z*_*i*_ from the ‘true values’ *x*_*i*_ of the quantity as follows:

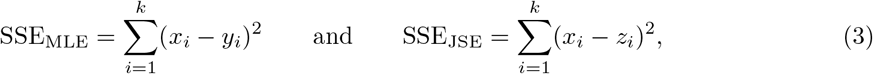

and compared these quantities.

Of course, the ‘true value’ of any quantity is unknowable except with infinite data. So, in the following, given long-enough trial durations for each participant, we use the parameter estimate using the full trial duration as the ‘true’ value, as a proxy for the real true value) – analogous to the baseball example (Fig. 1a). Also analogous to the baseball example, we use the estimates for some shorter time duration as the maximum likelihood estimate (MLE), to distinguish it from the ‘true’ value. Because the MLEs use shorter-than-available data, they are necessarily imperfect and the goal is to see if JSEs systematically improve their accuracy and move them toward the ‘true’ value. We will drop the quotes from ‘true’ in the following sections.

The computational cost of obtaining the JSE are low: once the MLEs *y*_*i*_, shrinkage factor *c*, and the MLE mean 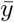 are known, equation 1 implies just three elementary arithmetic operations per JSE, so a total of 3*k* arithmetic operations; of course, computing the mean MLE and the shrinkage factor also require *O*(*k*) operations. So, overall, the JSE requires *O*(*k*) arithmetic operations: that is, the computational complexity grows linearly in the number of participants *k*.

### 2.2 Biomechanical datasets considered and quantities estimated

#### 2.2.1 Foot placement dynamics in level-ground treadmill walking

The experiments and the quantities estimated follow that in [5]. Eight participants (*N* = 8, 4 males and 4 females, with height 1.70 ± 0.09 m, age 31.88 ± 10.3 years and mass 67.15 ± 19.2 kg (mean ± s.d.)) walked on a treadmill at belt speed 1.3 m/s and zero incline for 2 minutes (Fig. 2a: left). Three-dimensional marker-based motion capture (Vicon T20, 100 Hz) was used to track the position of a sacral marker on the pelvis representing the body’s center of mass, and two heel markers representing each foot, used in the analysis (Fig. 2a: right).

**Fig. 2.**
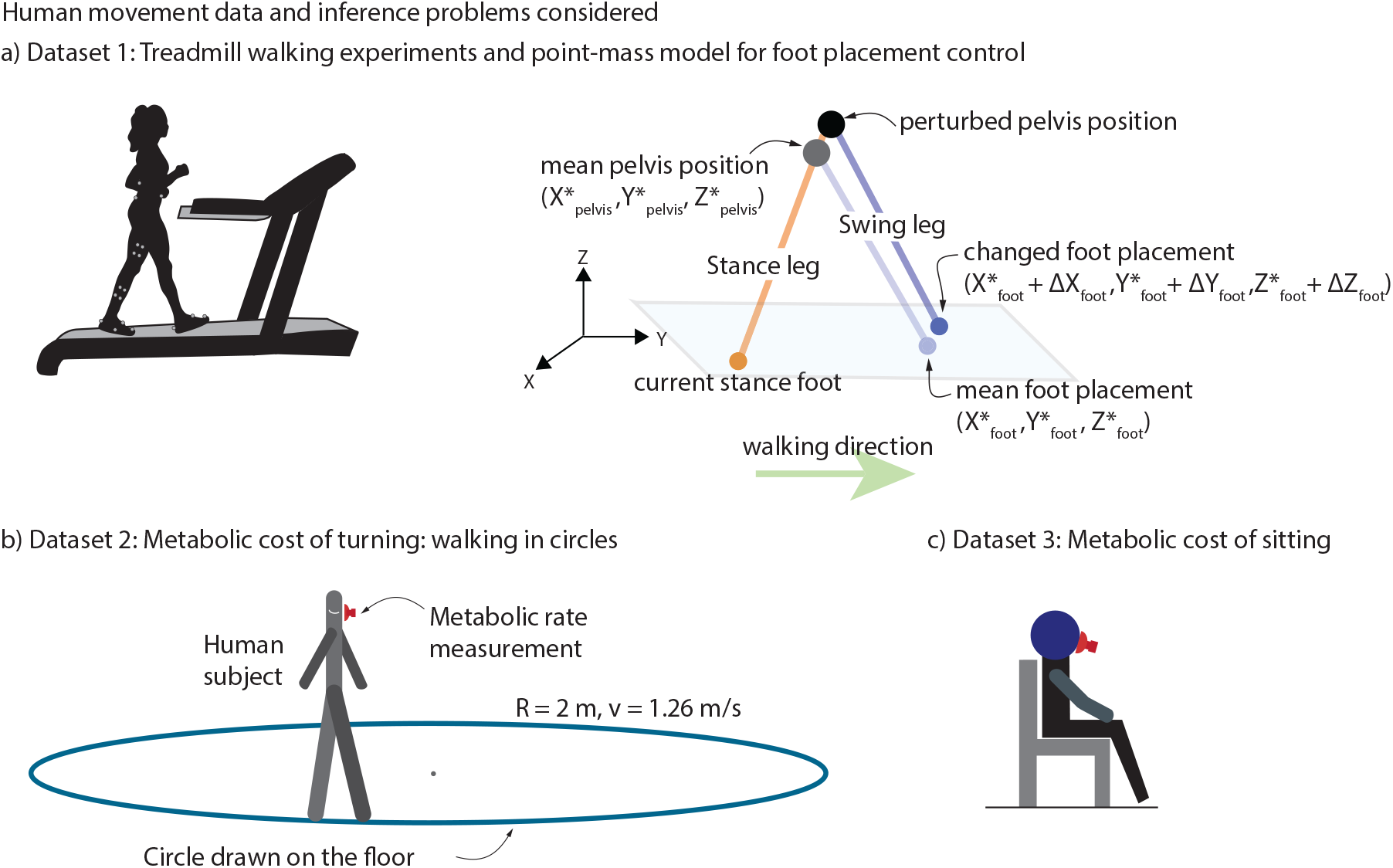
Biomechanical datasets used and quantities estimated. a) Treadmill walking experiments. Participant walks on a constant speed and level-ground treadmill with motion capture markers placed on pelvis, thighs, shanks, and feet. b) Point-mass model for foot placement control. Point-mass walking model with the human body represented by the position of pelvis as the center of mass and two massless legs. Foot placement in the next stance foot in response to deviation from the mean pelvis trajectory is also shown. c) Metabolic rate of turning: walking in circles. Participant walking on circle drawn on the floor of radius 2 m and walking at a 1.26 m/s speed while collecting metabolic rate measurements using a face mask. d) Metabolic rate of resting. Participant sitting in place while collecting metabolic rate measurements using a face mask.

Humans step in the direction of the perturbation to the center of mass [4, 5, 44, 45]. As in [4, 5, 44, 45], we estimate the change in foot placement following a deviation in the pelvis position (Fig. 2a: right). The motion data were analyzed by fitting linear models between deviations in pelvis states at mid-stance as input and the next foot position as output. The pelvis state at midstance *P* comprises the sideways position, sideways velocity, and forward velocity, that is, 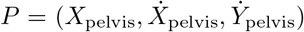. The next stance foot position is *S* = (*X*_foot_, *Y*_foot_). We used least squares regression to estimated the 2 × 3 Jacobian matrix (*J*) relating the next stance foot position to the midstance pelvis state as in the following linear model:

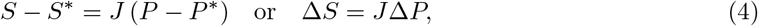

where *P* ^*∗*^ and *S*^*∗*^ are trial-wise mean values of *P* and *S* and Δ*P* and Δ*S* are deviations from these trial-wise means. The elements of the matrix *J* are sensitivities or partial derivatives of the individual output variables relative to the individual input variables: for instance, *J* (1, 1) is the sensitivity of sideways foot placement (*X*_foot_) to deviations in sideways pelvis marker velocity 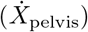 at mid-stance and can be denoted as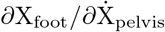. See [4, 5, 44, 45] for more details.

For illustrative purposes, we applied the JSE only to one element of the *J* matrix, namely the sensitivity 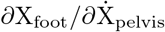. Each trial had a minimum of 100 steps. So for each participant, we computed the true value of the partial derivative 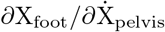 using the 100 steps, the maximum available data. We also computed this quantity for a series of smaller trial durations ranging from 15 to 90 steps using the same procedure. The error covariance 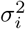 of these estimates are obtained directly from the least squares regression for each participant-*i* and each trial duration.

### 2.2.2 Metabolic rate during walking in a circle

The experimental data for this section is drawn from [18], with *N* = 11 participants: 6 males and 5 females, with height 1.74 ± 0.10 m, age 33.55 ± 22.2 years and mass 74.01± 12.5 kg (mean ± s.d.). Participants walked along marked circles on the ground at four different circle radii (*R* = 1 m, 2 m, 3 m, 4 m) and four different constant tangential speeds ranging from 1.58 m/s to 1.8 m/s for a total of 16 trials (Fig. 2b). Participants walked for six to seven minutes to achieve a sufficient metabolic steady state. Respiratory oxygen and carbon dioxide flux was measured using indirect calorimetry (Oxycon Mobile). For illustrative purposes, we only considered the trials with radius 2 m and approximate speed of 1.26 m/s.

We computed metabolic rate per unit mass, *Ė*, using the Brockway equation [15]:

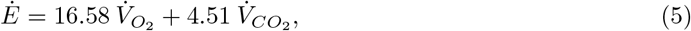

where *Ė* is in W*/*kg and 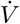 is in mL s^*−*1^ kg^*−*1^. To estimated the steady state from any fraction of the total trial duration (e.g., first 2 minutes), we fit an exponential *Ė* = *a*_0_ + *a*_1_*e*^*−t/τ*^, where *τ* is the time-constant.

Each participant’s trial had a minimum of 54 data points. So, for each participant, we compute the true value to be the steady state metabolic rate *Ė* computed from the 54 data points using the exponential fit. Similarly, we compute the steady state metabolic rate *Ė* using the exponential fit for shorter duration fractions of the trials, ranging from 15 data points (about 25 seconds) to 51 points (about 2 minutes). For such metabolic data, the number of data points is correlated with the trial duration but the proportionality is not perfect because of breathing rate variability (see appendix A and Fig. A1). Error variance 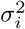 of the individual steady state estimates are computed using bootstrap statistics.

### 2.2.3 Metabolic rate during resting

Analogous to walking metabolic energy expenditure analysis from subsection 2.2.2, we compute *Ė* for quiet sitting experiments. We collected these energy expenditure datasets from multiple previously conducted experiments [17, 18, 42, 43], giving data for *N* = 27 participants: 23 males and 4 females, with height 1.79 ± 0.05 m, age 24.41 ± 4.5 years and mass 75.42± 9.9 kg (mean ± s.d.). In these studies, as a baseline for metabolic rate of movement, participants were asked to sit quietly for 6-7 minutes and measure oxygen *O*_2_ and carbon dioxide *CO*_2_ volume rates to estimate resting metabolic rates via a simple average over the duration.

Each participant’s resting trial had at least 40 data points, so we compute the ‘true value’ to be the mean metabolic rate *Ė* over all the 40 data points. We also computed the mean for shorter fractions of the trial duration, ranging from 10 data points to 35 data points; these estimates are labeled ‘MLEs’. Error variance *σ*^2^ for the individual estimates are computed by first computing the data deviations from mean for each participant, pooling these deviations over all participants, and determining the standard error *σ* from these deviations via bootstrap.

## 3 Results

### The JSE can move data towards the truth

Fig. 3 shows how JSE improves the MLEs obtained from a particular short duration trial for each participant, moving the individual MLE’s toward the mean of all the MLEs. This results in the spread of the JSEs being lower than the spread of the MLEs, and closer on average to the ‘true’ values obtained from the longer duration trials in terms of summed squared errors.

**Fig. 3.**
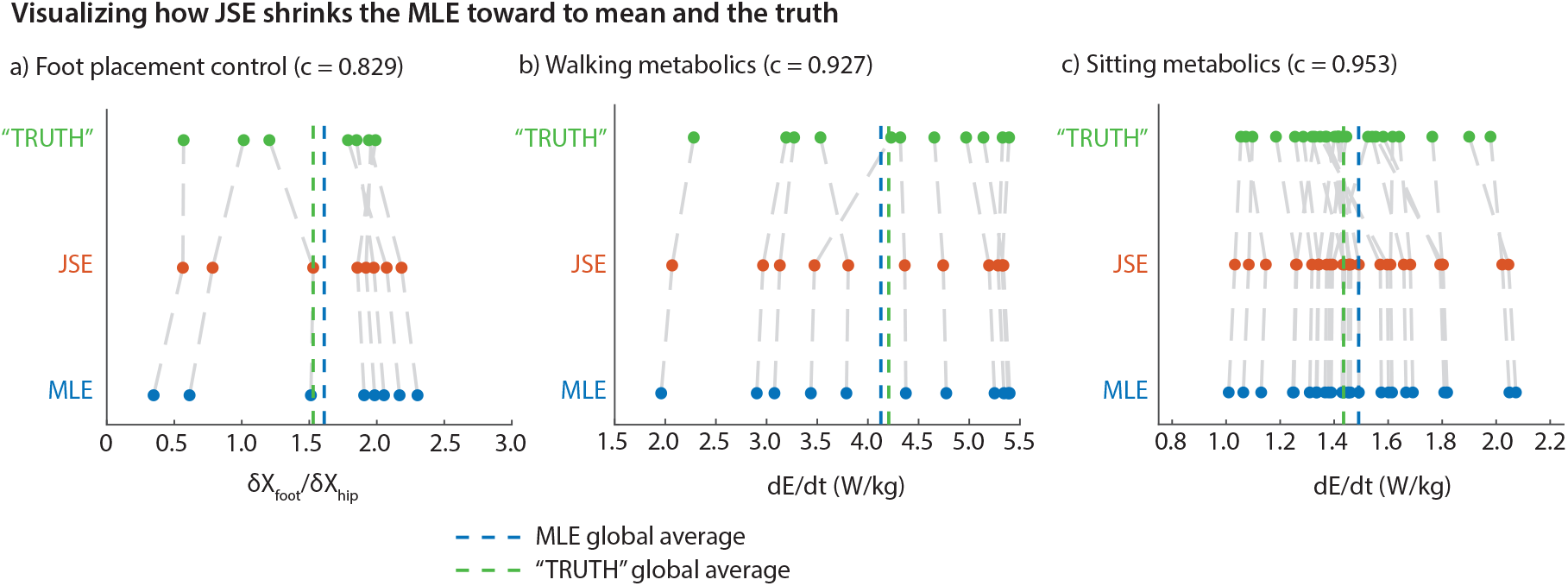
Visualizing how JSE shrinks the MLE toward (a) Walking kinematics: model comparison between the JSE and the MLE relative to the TRUTH for coefficient *δX*_Foot_*/δX*_Hip_ using 25 steps or data points of MLE data from 100 steps or data points of TRUTH data. Number of participants, *k*, is 8 and they correspond to the scatter points. (b) Walking metabolics: model comparison between the JSE and MLE relative to the TRUTH for *Ė* using 21 data points of MLE data from 54 data points of TRUTH data. Number of participants, *k*, is 11 and they correspond to the scatter points. (c) Sitting metabolics: model comparison between the JSE and MLE relative to the TRUTH for *Ė* using 20 data points of MLE data from 40 data points of TRUTH data. Number of participants, *k*, is 27 and they correspond to the scatter points. Each data point in the scatter plot represents a participant’s individual estimate of a kinematic or a metabolic parameter using full or subset of full trials.

For foot placement gain *∂X*_Foot_*/∂X*_Hip_, when the MLEs were obtained from only 25 steps, the JSEs shrunk the MLEs by about 17% towards their mean (*c* = 0.829, Fig. 3a, d). This resulted in a summed squared error for the JSEs about 46.9% lower than that of MLEs (*p* = 0.0475, paired *t*-test). For circle walking metabolic rate, when the MLEs were obtained using only 21 data points, the JSEs shrunk the MLEs toward their mean by about 7% (*c* = 0.927, Fig. 3b, e). This resulted in a summed squared error for the JSEs about 26.3% lower than that for the MLEs (*p* = 0.0272, paired *t*-test). Finally, for resting metabolic rate, when the MLEs were obtained using only 20 data points, the JSEs shrunk the MLEs by 5% toward their mean (or c = 0.953). This resulted in a summed squared error for the JSEs about 9.7% lower than that for the MLEs (*p* = 0.0149, paired *t*-test).

While the SSE with respect to the true values is reduced by the JSEs, we generally do not expect the means of the true value, MLEs, and JSEs to be significantly different. That the means of MLE and the true values are close together and unbiased is why JSE is defined as moving the MLEs toward their grand mean. Performing paired *t*-tests, we find that means of true values and MLEs are not significantly different for foot placement control (*p* = 0.508) and not significantly different for walking metabolic rates (*p* = 0.573). However, true and MLE values were significantly different for the resting metabolic rates (*p* = 0.009), suggesting that the MLE sample is slightly biased from the true sample, potentially explaining the lower error reduction by JSE for the resting metabolic rates. Of course, by definition, the JSEs and MLEs have the same mean, and *t*-tests confirm that they are not significantly different (*p* = 1).

### JSE reduces error more for shorter duration trials

For the range of short duration trials we considered, we found that the JSE increased accuracy compared with MLE in all three estimation problems (Fig. 4). Specifically, when compared with MLE, the summed squared error (SSE) for the JSE was lower by about 13% to 56% for foot placement control, 4% to 85% for circle walking metabolic rate, and 5% to 14% for resting metabolic rate data, with the higher reductions being for the shorter duration trials (Fig. 4a-f). Both the absolute reduction in error (Fig. 4a-c) and the relative reduction in error (Fig. 4d-f) generally decrease as the MLE sample size approaches the ‘true’ sample size. Correspondingly, the shrinkage factor *c* increases with increasing MLE sample size for all estimation problems, approaching *c* = 1.0, indicating that shrinkage decreases with larger MLE sample sizes. The most shrinkage (low *c* values) occurs at small sample sizes, with foot placement kinematics and walking metabolics experiencing relatively higher shrinkage than sitting metabolics estimation problem.

**Fig. 4.**
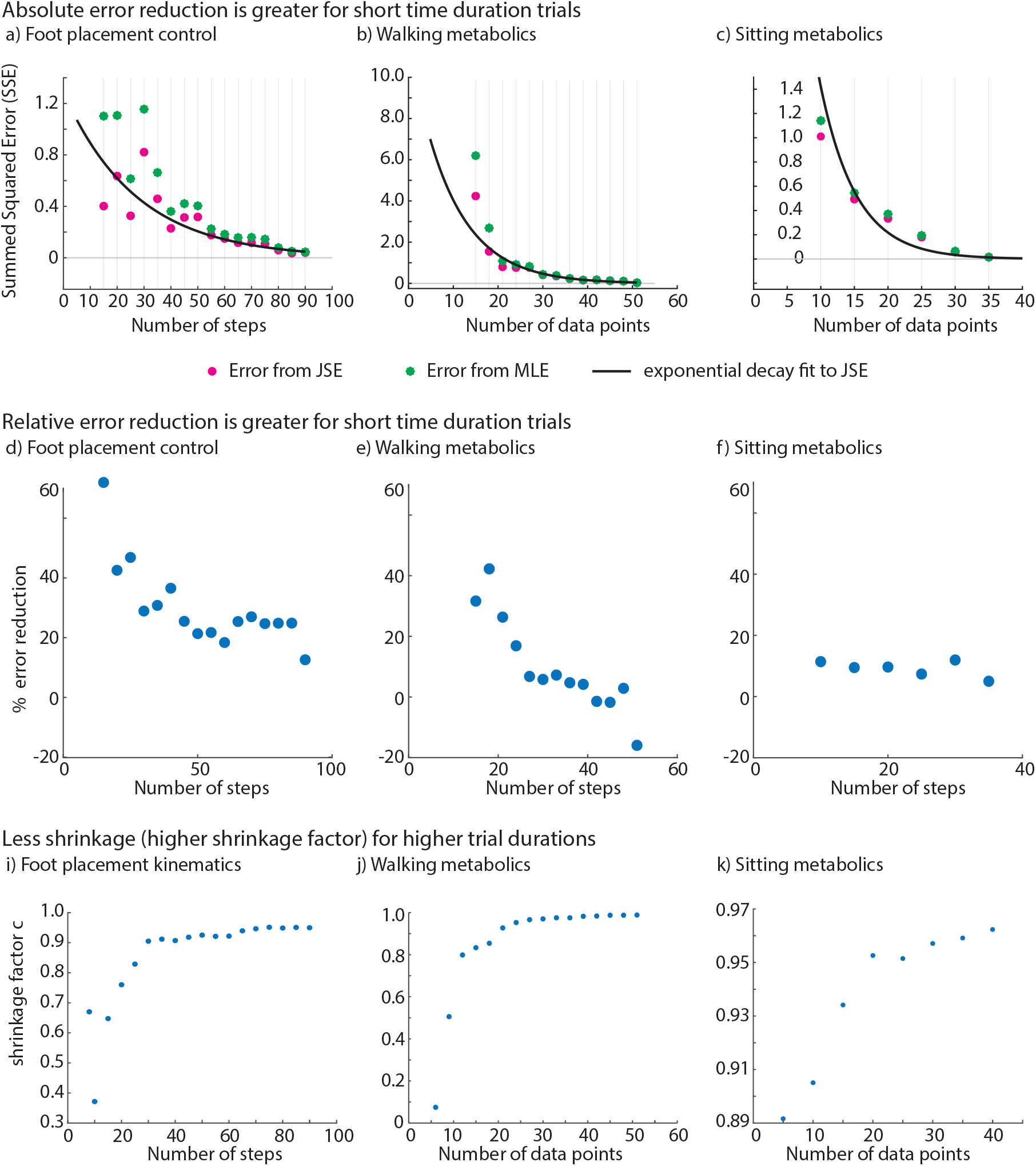
The James-Stein estimator (JSE) led to reduction in summed squared errors (SSE) compared with maximum likelihood estimation (MLE) for foot placement kinematics, walking energy rate and resting energy rate. **a), b), c)** Summed squared error (SSE) values for the MLE and the JSE datasets for each parameter estimation problem. Bestfit curve is overlaid on the scatter points for the error from JSE shown in pink color. We fit the SSE from JSE to an exponential decay model of the form: 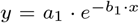 where the parameters *a*_1_ and *b*_1_ are estimated using linear regression on the log-transformed SSE from JSE data, and *x* is the number of steps or data points. **d), e), f)** Percent reduction in summed squared error for each parameter estimation problem. **i), j), k)** Shrinkage factor *c* for each parameter estimation problem.

When the MLE sample size is very close to the true sample size, the JSE sometimes ‘apparently’ performing worse relative to MLE, as illustrated in the case of walking metabolics, where we get negative percent reduction in error (Fig. 4e). This artifact can be explained by observing that because the true sample size is finite, so what we call the true estimate already contains some random error. Thus, when the MLE sample size gets too close to the true sample size, the error estimate for MLE and JSE become unreliable because of comparable error in the ‘true’ value. Of course, in this limit, we might as well use the full trial duration estimate, as the improvement due to JSE’s improvement is quite small anyway and the trial durations for the MLE and the ‘true’ estimate are not that different. Alternatively, we could apply JSE to the full duration estimate to obtain a further slight improvement.

At the other extreme, when the sample size is too small — smaller than what we have considered in Fig. 4 – the MLE has high error and the JSE may be even more unreliable due to poor estimation of the within participant error *σ*_*i*_ in the JSE formula (equation 1). So, it is not advisable to use such small sample sizes.

## 4 Discussion

We introduced the James-Stein estimator (JSE) as an approach to enhance the accuracy of parameter estimation in the context of biomechanical data, particularly when dealing with limited sample sizes. We applied the JSE to three different time-series datasets: foot placement control during walking, steady-state metabolic energy rate during walking, and mean resting metabolic energy rate during sitting. We observed that the JSE effectively shrinks individual maximum likelihood estimates (MLEs) towards the global average, bringing them closer to the true values as a whole. This shrinkage improved the estimation accuracy, reducing the summed squared error (SSE). Overall, the JSE demonstrated its potential to enhance parameter estimation in biomechanical studies, especially when working with smaller sample sizes and larger variability.

While the JSE reduced SSE in all estimation problems, it is more effective when the datasets are such that the individual estimates have lower accuracy compared with between-participant variance. These properties may arise from inter-participant variability, measurement noise, or sample size limitation. The JSE was successful at substantially reducing SSE in the case of foot placement kinematics and walking metabolics estimation problems, but only a more minimal reduction in MSE was obtained for the observed in the resting metabolic rate dataset, where the within-individual variance was relatively lower.

The theorems guaranteeing JSE outperforming MLE by minimizing SSE were originally derived under the following assumptions: the dimension of the quantities being estimated is greater than or equal to three, the variables are independent from each other, and the underlying distribution is normal [8, 10, 25]. However, the normality assumption is not critical to JSE and violating it only reduces the accuracy improvements obtained [10]. The additional error is negligible if the number of quantities estimated, *k*, is greater than 15 and minimal if *k* is as low as 9 [10]. In the context of biomechanical data, if we estimate one quantity per human participant from their respective time-series data, the number of quantities estimated, *k* equals the number of individuals. So we expect greater insensitivity to assumptions when there are more individuals.

The shrinkage factor *c* as computed from equation 2 may not always be positive. If *c* is negative, the James-Stein estimator moves the estimate ‘beyond’ the global average and the performance of JSE is compromised. In these cases, the positive-part JSE *z*^+^ should be implemented [46]: that is, when the shrinkage factor *c* is negative, replace it by zero, avoiding error increase due to JSE. The positive-part JSE dominates the regular JSE, that is, better than the regular JSE in some cases, while being identical to it in other cases [46].

James-Stein estimation can be related to and contrasted with Bayesian inference [47]: in classical Bayesian inference, there is a fixed ‘prior’ distribution and a new data point is corrected toward the mean of this prior distribution (Fig. 5a); in James-Stein estimation, the JSEs are obtained by reducing the larger MLE values and increasing the smaller MLE values, bringing them closer to the grand MLE average (Fig 5b) – and this can be interpreted as moving the MLEs toward an empirically-based prior obtained from the individual MLEs (Fig 5c). If reliable prior distributions are known for the parameters of interest, it may be better to use a classical Bayesian approach in lieu of JSE, whereas JSE has the advantage of computational simplicity and not needing a prior. Given JSE’s interpretation as an empirical Bayes estimator with simple assumptions [25, 47], an intermediate approach between JSE and a classical Bayesian approach could be an empirical Bayes approach that more precisely estimates an empirical prior before the posterior is computed [27]. Alternatively, when other structural information is available about the parameters, such information can be used to improve the statistical estimates: for instance, researchers have used isotonic regression when it is known that the parameters to be estimated are ordered with a known order [30]; similarly, if it is known that most of the parameters to be estimated have zero values, one could use a pretest estimation procedure [30]. Our work involved estimating parameters that did not have such known properties and reliable prior distributions of these parameters are not available.

**Fig. 5.**
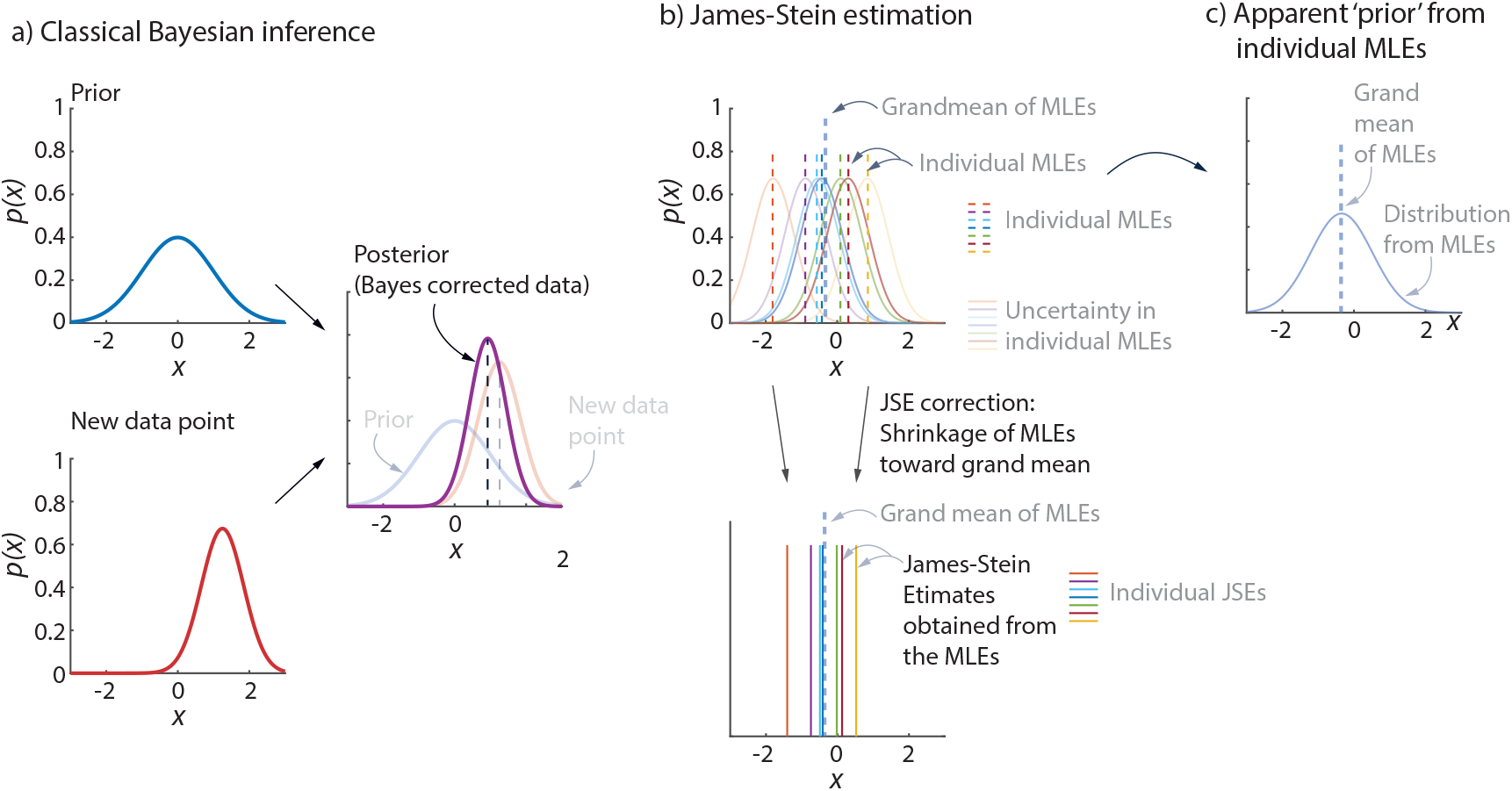
James-Stein estimation and Bayesian inference. a) Classical Bayesian inference uses a fixed ‘prior’ and a new data point with its own error distribution (‘likelihood’) to get a better estimate, namely the ‘posterior.’ b) Given MLEs for a few participants, with their corresponding error distributions, James-Stein estimation simply shrinks the MLEs toward the grand mean of all the MLEs. c) One can interpret the JSEs as being related to an empirical Bayes correction, in which a prior is implicitly obtained from all the MLEs.

By lowering the summed squared error for a given trial duration (Fig. 4), JSE can potentially allow using shorter trial duration for a given accuracy. If a particular typical error value is sought, one can use a plot like Fig. 4 to read off the corresponding trial duration that would provide this desired accuracy level when using JSE.

By improving estimation accuracy and efficiency on average, JSE has the potential to provide better biomechanical personlization. While JSE decreases overall mean squared error on average, some participants may experience worsened outcomes, reflecting that the JSE does not provide individually better estimates for every parameter [48]. We showed statistically significant improvement in the accuracy for all three datasets we examined. However, such improvements are only for some range of sample sizes and clearly not for all individuals. Thus, using JSE in lieu of MLE requires a value judgement to improve the overall error across participants, while potentially making estimates of some participants worse — especially outlier individuals. This tradeoff is analogous to similar value judgment in other medical decision-making [49, 50]. For example, different patients may have varying responses for a fixed drug dose, which could be established through clinical trials, potentially suggesting individualized medicine approaches. However, a truly individualized prescription may not always be practical. So the next best solution is to account for inter-participant variability in drug response via demographics, anatomical variations, etc., thus implicitly improving typical individual outcome by studying population-level variability – potentially at the cost of affecting outlier individuals adversely. To implement JSE in a biomedical or wearable device design context, one would process patients or participants in groups — or potentially use the history of patient or participant data or perhaps their summary to improve the device or estimate for the most recent patient.

The original JSE formula [25, 28] was derived by considering that the individual means (MLEs) were drawn from a Gaussian distribution and are independent. So, while high variance in these means (representing one kind of data heterogeneity) is naturally accommodated by JSE, larger heterogeneity in the dataset – for instance, consisting of multiple distinct participant populations, each drawn from a different Gaussian with a distinct mean – can degrade performance of JSEs. In such cases, it may be meaningful to cluster the data by other relevant participant characteristics before applying JSEs to the individual clusters.

One might ask the following question: ‘if the JSE is indeed better than the MLE in terms of accuracy, why has it not replaced the MLE yet?’ [51]. Efron [51] partially answers this question by stating that the statistical community often adhere to conventional methods such as the MLE to protect individual inferences from the influence of group-based approaches [51]. In addition, researchers often prefer the familiarity, simplicity, and widespread applicability of the MLE compared with other estimators. Finally, the MLE is often preferred because it has zero bias, that is, it does not tend to underestimate or overestimate a population parameter. JSE increases bias to reduce variance and mean squared error [25, 52–54] (Appendix B and Fig. B2). So a user may need to decide how much bias is acceptable while implementing the JSE.

In summary, James-Stein estimator could be a valuable tool for improving parameter estimation in biomechanical studies, particularly when sample sizes are small, or variability among participants is high. Future work could consider biomechanical data of populations with limited mobility (i.e. short and variable) to improve accuracy for such datasets. Additionally, introducing JSE to other types of datasets, such as high-dimensional biomechanical data from wearable sensors, could also open up new possibilities for enhancing prediction accuracy in personalized healthcare applications.

## Acknowledgments

The work was supported in part by NIH grant R01GM135923-01 and NSF SCH grant 2014506.

## Data availability

The data for this manuscript is available at http://datadryad.org/stash/share/fVGgnWsMzPAxO1gPrGPuOXEAm0msSDd79KxjmF7qVAo for review purposes and will also be available publicly on the Dryad database at DOI:10.5061/dryad.3j9kd51v9.

## Appendix A Trial durations for metabolic data

For a given number of data points, trial duration is not exactly but only approximately consistent across participants. This is because Oxycon system collects data breath by breath and does not have a fixed sampling rate. e.g., a 2-minute trial may consist of 35 up to 60 data points (Fig. A1).

**Fig. A1.**
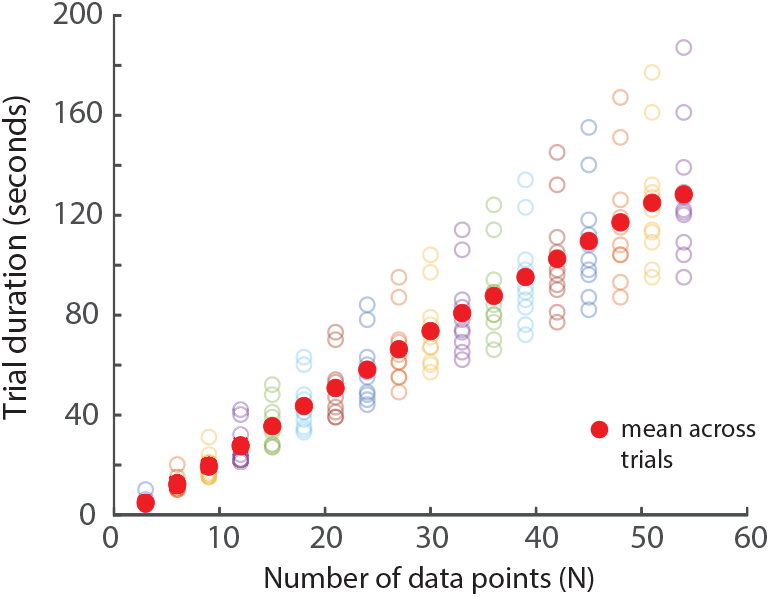
Trial durations of all participants circle-walking metabolic trials in seconds. The mean values are indicated by bolded red circles. Variations in trial duration among participants result from differences in their breathing rates.

## Appendix B James-Stein estimator optimizes bias-variance trade-off

The James-Stein estimator can be understood as minimizing the mean squared error given all the data, by optimally trading off between bias (systematic error) and variance (random error). We can estimate the bias and variance (*σ*^2^) as follows, with the Mean Squared Error (MSE) being the sum of these two quantities [25, 53, 54]:

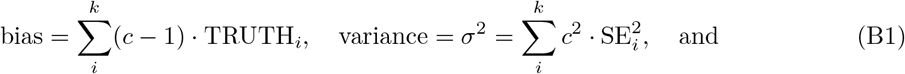

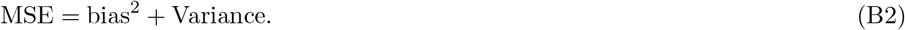

The JSE decreases the overall MSE by reducing the variance at the cost of increasing the bias: see Fig. B2 for an illustration using the foot placement walking dataset [8, 25].

**Fig. B2.**
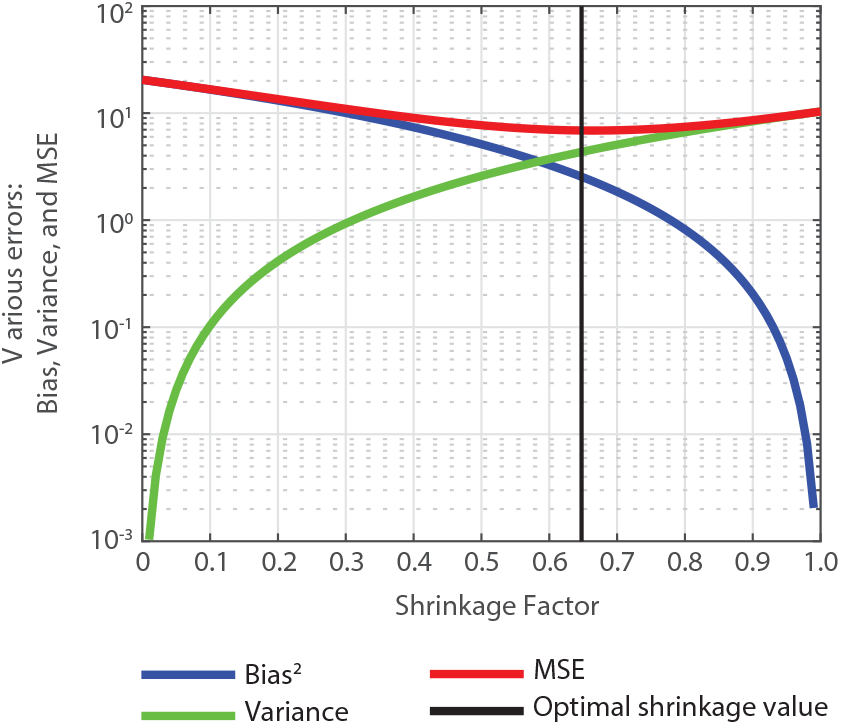
Bias, variance, and Mean Squared Error (MSE) as a function of the shrinkage factor *c*, for a representative MLE sample size of 10 steps from the foot placement walking data set. The optimal computed shrinkage value *c* for this is about 0.648, shown as a black vertical line.

